# Exposure to HIV alters the composition of maternal microchimeric T cells in infants

**DOI:** 10.1101/2024.03.01.583002

**Authors:** Blair Armistead, M. Quinn Peters, John Houck, Marc Carlson, Christina Balle, Nolawit Mulugeta, Clive M. Gray, Heather B. Jaspan, Whitney E. Harrington

**Affiliations:** Center for Global Infectious Disease Research, Seattle Children’s Research Institute, Seattle, Washington, USA; Research Scientific Computing, Enterprise Analytics, Seattle Children’s Research Institute, Seattle, Washington, USA; Division of Immunology, Department of Pathology, Institute of Infectious Disease and Molecular Medicine, University of Cape Town, Cape Town, South Africa; Department of Global Health, University of Washington, Seattle, Washington, USA; Department of Pediatrics, University of Washington, Seattle, Washington, USA; Division of Immunology, Biomedical Research Institute, Stellenbosch University, Cape Town, South Africa

**Author notes:** Corresponding author: Dr. Whitney Harrington,.

## Abstract

Infants exposed to HIV but uninfected (iHEU) display altered cellular immunity and are at increased risk of infection through poorly understood mechanisms. We previously reported that iHEU have lower levels of maternal microchimerism (MMc), maternal cells transferred to the offspring in utero/during breastfeeding. We evaluated MMc levels in T cell subsets in iHEU and HIV unexposed infants (iHU) to determine whether a selective deficiency in MMc may contribute to altered cellular immunity. Across all infants, MMc levels were highest in CD8+ T cells; however, the level of MMc in the CD8 T cell subset was significantly lower in iHEU compared to iHU.

## Background

Large-scale implementation of programs to prevent maternal-to-child transmission of HIV-1 have substantially reduced the incidence of HIV infection in infants (1). Still, each year, nearly 1.3 million infants are born to mothers living with HIV but are themselves uninfected (iHEU) (1). Compared to infants who were unexposed to HIV in utero or during breastfeeding (iHU), iHEU experience greater morbidity and mortality associated with infectious disease including diarrheal disease and respiratory infection (2, 3), which are not completely explained by factors such as maternal health, breastfeeding, or socioeconomic status (3, 4). Several studies have demonstrated altered CD4+ and CD8+ T cell responses in iHEU following pathogenic challenge or vaccination (5, 6). However, the mechanisms underlying differential T cell immunity in iHEU are poorly understood.

We recently reported that iHEU had significantly lower levels of maternal microchimerism (MMc) at birth compared to iHU, and that lower MMc at birth was associated with attenuated response to BCG vaccination (7). MMc refers to the transmission of maternal cells to offspring *in utero* or during breastfeeding. MMc cells are acquired as early as the second trimester of pregnancy (8, 9) and can persist for decades after birth (10). In cord blood mononuclear cells from healthy newborns in the United States, the greatest levels of MMc were found in memory T cells (11). In addition, recent work in mice suggests a role of MMc T cells in fetal and pup immune development and protection against infectious challenge (12, 13, 14). We hypothesized that a selective deficiency in MMc within T cell subsets might contribute to altered cellular immunity in iHEU. In a cohort of South Africa infants, we identified a higher level of MMc in CD8+ T cells versus CD4+ T cells. However, the level of MMc within CD8+ T cells was lower in iHEU versus iHU, highlighting a potential mechanism underlying altered cellular immune responses in these infants.

## Methods

### Cohort

Samples and data were utilized from the Innate Factors Associated with Nursing Transmission (InFANT) study, an ongoing prospective cohort study of mothers and their infants conducted in Khayelitsha, Cape Town, South Africa (7) Mothers and their infants included here were enrolled upon delivery from March 2014 to March 2018. Voluntary counseling and HIV testing was performed at the time of antenatal care registration. Pregnant women living with or without HIV were eligible for inclusion. Exclusion criteria included delivery before 36 weeks of gestation, birthweight lower than 2.4 kg, and major pregnancy or delivery complications. All infants classified as iHEU were confirmed HIV negative by polymerase chain reaction (PCR) at delivery and at later timepoints. After birth, infant follow-up visits occurred at weeks 7, 15, and 36 of life. For the present study, samples were included for infants where we had previously identified MMc in whole blood from that timepoint (7) and a where a matched peripheral blood mononuclear cell (PBMC) sample was available (n=19). Two of the infants contributed samples from two different timepoints.

### Isolation, storage, and thawing of peripheral blood mononuclear cells

PBMC were isolated from infant whole blood using ficoll centrifugation. PBMC were resuspended in 90% fetal calf serum with 10% dimethyl sulfoxide, and aliquoted into cryovials for storage in liquid nitrogen. Prior to experiments, PBMC were thawed at 37ºC, added dropwise to pre-warmed Rosewell Park Memorial Institute (RPMI) medium with L-glutamine and 20% fetal bovine serum, and centrifuged. Resulting cell pellets were resuspended in complete medium (RPMI-1640 with L-glutamine, 10% FBS, 100 U/mL penicillin, 100μg/mL streptomycin), counted, and assessed for viability.

### Magnetic activated cell sorting and MMc quantitation of infant immune cell subsets

To separate infant PBMC into CD8+, CD4+, and non-CD8/CD4 T cell fractions, sequential magnetic activated cell sorting (MACS) was performed. First, cells were strained using a 100μm nylon mesh filter, and CD8+ T cells were isolated using the REAlease CD8 T Cell MicroBead Kit (Miltenyi) according to manufacturer instructions. CD4+ T cells were subsequently isolated from the resulting CD8-eluent using the REAlease CD4 T Cell MicroBead Kit (Miltenyi) according to manufacturer instructions, generating a fraction of CD4+ T cells and a fraction of CD4-/CD8-cells, which were designated as “other” immune cells. During initial protocol development, purity of the CD8+ T cell and CD4+ T cell fractions were found to be ∼90% by flow cytometry. Following sequential MACS separation, genomic (gDNA) was isolated from each immune cell fraction with the Prepfiler Forensic DNA Extraction kit (ThermoFisher) per manufacturer instructions.

### MMc quantitation in cellular subsets

Human leukocyte antigen (HLA) and non-HLA typing had been previously conducted for all mothers and infants to identify a maternal-specific allele for each pair, as described elsewhere (7). The maternal-specific allele identified for each maternal-infant pair was selectively amplified from gDNA in each sorted cell population using a panel of previously developed quantitate PCR (qPCR) assays (11). Briefly, gDNA were divided into multiple reaction wells to exhaust the substrate. A calibration curve for the polymorphism-specific assay was included to quantify the level of MMc in each reaction well, and the MMc genomic equivalent (gEq) was summed across wells. To quantify the total gEq tested in each assay, qPCR targeting the β-globin gene was performed for each sample and evaluated alongside a β-globin calibration curve (human gDNA, cat# G1521, Promega).

### Statistical analyses

The frequency of MMc across CD4+ T cells, CD8+ T cells, and the non-CD4+/CD8+ subset was assessed with negative binomial regression, which accounts for the number of microchimeric gEq and the total gEq tested in each sample with clustering by individual (15). HIV exposure and infant age were associated with overall MMc level in primary analyses and were included as covariates in the model. In addition, there was a significant interaction between HIV exposure and MMc level in T cell subsets (p <0.1), and thus, we also present HIV exposure stratified models.

## Results

### Cohort characteristics

We included 19 samples obtained at birth (n=3, 15.8%), week 15 (n=12, 63.2%), and/or week 36 (n=4, 21.1%) of age from 17 infants. Just under half of the mothers were living with HIV (8, 47%); of these mothers, 5 (62.5%) had initiated antiretroviral therapy (ART) prior to the current pregnancy, and 3 (37.5%) initiated ART during pregnancy (Table 1). Eleven (64.7%) of the infants were breastfed at the time of sample collection.

**Table 1.** Study population characteristics.

|  | Total (n=17)* |
| --- | --- |
| **Infant sex, female, n (%)** | 10 (58.8) |
| **Gestational age at delivery, weeks, mean (SD)** | 39 (1.6) |
| **Age at time of sampling, n (%)\*** |  |
| Birth (0 weeks) | 3 (15.8) |
| 15 weeks | 12 (63.2) |
| 36 weeks | 4 (21.1) |
| **HIV exposed, n (%)** | 8 (47.1) |
| **Receiving breastmilk, n (%)** | 11 (64.7) |

\*n=17 infants contributed a total of 19 samples
SD, standard deviation

### MMc is enriched in CD8+ T cells versus CD4+ T cells but diminished in iHEU

In the primary analyses, we tested the individual effect of HIV exposure, infant sex, and breastfeeding on MMc across all sorted cell subsets. HIV exposure was associated with lower MMc (IRR: 0.04, p=0.004), but there was no association between breastfeeding or infant sex and MMc level across all subsets. We next compared the frequency of MMc in each sorted subset and found greater levels of MMc in CD8+ T cells compared to CD4+ T cells (incident rate ratio (IRR): 43.9, p<0.0001) and to other immune cells (IRR: 35.7, p<0.0001) (**Fig. 1A**). We then tested whether HIV exposure was associated with MMc level within each sorted subset. iHEU had lower levels of MMc in CD8+ T cells (IRR: 0.035, p=0.001), but not CD4+ T cells (p=0.74) or other immune cells (p=0.15), relative to iHU (**Fig. 1B**). As a result, subsequent stratified analyses revealed that the difference in MMc frequency in CD8+ versus CD4+ T cells was substantially greater in iHU (IRR: 53.6, p<0.0001) compared to iHEU (IRR: 4.8, p=0.003) (**Fig. 1C**).

**Fig. 1.**
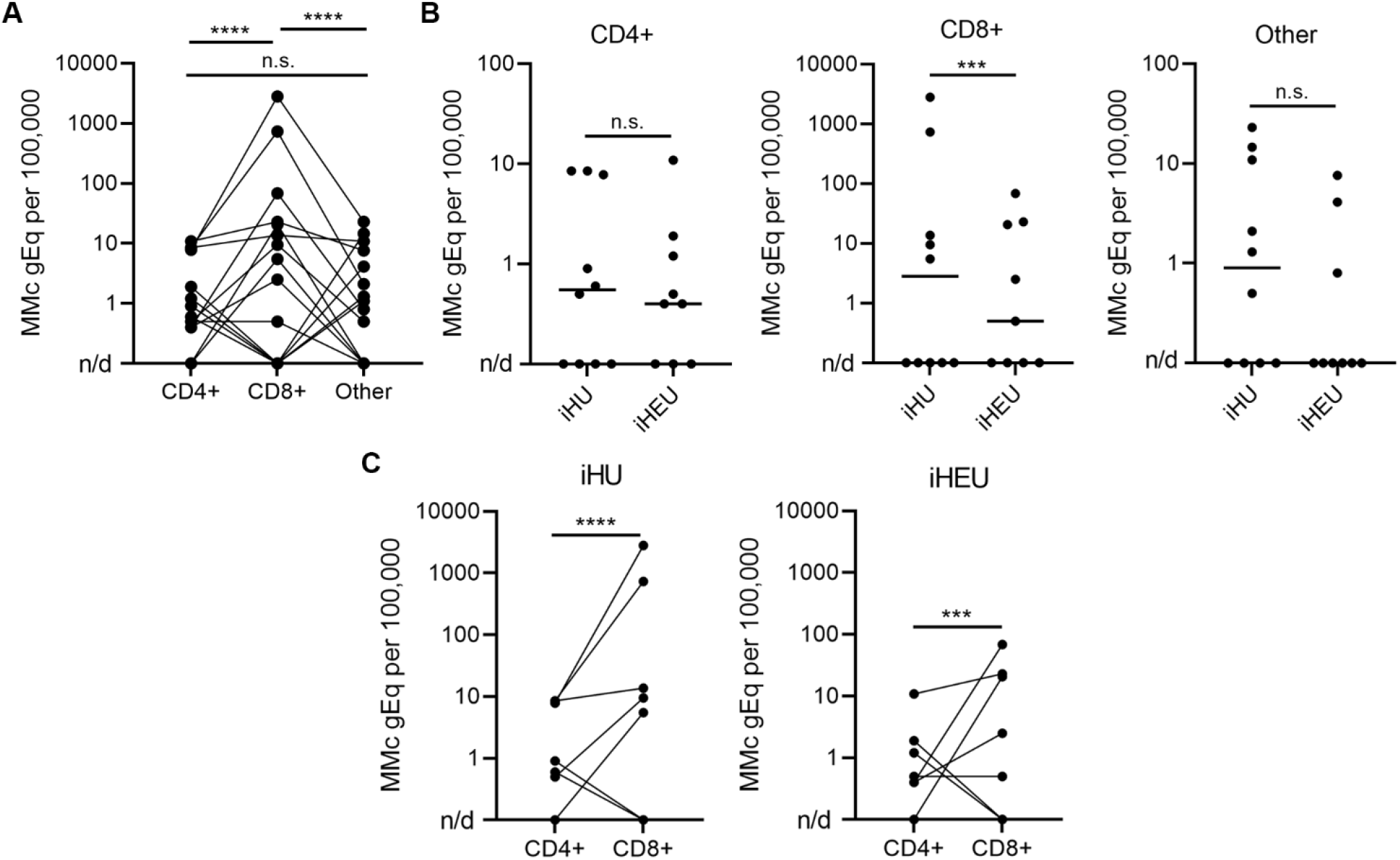
MMc is enriched in CD8+ T cells but this effect is diminished in iHEU. Infant PBMC (n=19) were sorted into CD4+ T cells, CD8+ T cells, and other cells using MACS, and MMc in each sorted subset was quantitated using qPCR targeting a maternal-specific allele. Comparisons of MMc level was conducted with negative binomial regression models accounting for the number of microchimeric gEq and the total gEq tested in each sample. **A**) The greatest levels of MMc were in the CD8+ T cell compartment. **B**) HIV exposure was associated with lower levels of MMc in CD8+ T cells but not CD4+ T cells or other immune cells. **C**) Enrichment of MMc in CD8+ T cells over CD4+ T cells was diminished in iHEU (IRR: 4.8) compared to iHU (IRR: 53.6). **** p<0.0001; *** p<0.001; n.s., not significant; n/d, not detected. gEq, genomic equivalent; iHEU, infant exposed to HIV but uninfected; iHU, infant unexposed to HIV; MACS, magnetic activated cell sorting; MMc, maternal microchimerism; PBMC, peripheral blood mononuclear cells; qPCR, quantitative PCR

## Discussion

In this study, we determined the level of MMc in sorted CD4+ T cells, CD8+ T cells, and other immune cells in PBMC from South African infants with and without HIV exposure. We report higher MMc levels in CD8+ T cells than CD4+ T cells and other immune cells across all infants, and HIV exposure was associated with a selective reduction in MMc CD8+ T cells. Our observations build upon the limited existing literature examining MMc in immune subsets in human cohorts, which report that memory T cells make up a substantial portion of the MMc cell graft inherited by infants (11). While one previous study evaluated fetal microchimerism (FMc) in maternal CD4+ T cell and CD8+ T cell subsets (16), to the best of our knowledge, our findings represent the first report of MMc in infant T cell lineage subsets.

The observed selective deficiency of CD8+ MMc T cells in iHEU may lend insight into mechanisms underlying differential cellular immune responses and increased susceptibility to viral respiratory and gastrointestinal infection in these infants. For instance, a reduction of maternally derived CD8+ T cells in iHEU could result in diminished viral control or decreased education of the infant immune response. In support of this notion, recent work in mice demonstrated that MMc T cells promoted the maturation of pup monocytes leading to augmented protection against viral challenge in the offspring (12). In addition, a recent study in mice demonstrated that *in utero* transmission of ovalbumin (OVA)-specific MMc T cells was associated with enhanced activation in offspring T cells and protection against challenge with OVA recombinant *Listeria* (13). Further work is needed to determine the role of MMc CD8+ T cells in cellular immunity in human infants, whether through indirect influence on fetal and infant immune development or through direct action of antigen-specific T cells.

Our study had several limitations. The total number of PBMC obtained from infants was low, which precluded our ability to sort into additional T cell populations, such as memory and naïve subsets. In addition, for most infants, we measured MMc at a single timepoint. Therefore, we are unable to determine whether the enrichment of MMc in CD8+ T cells over CD4+ T cells persists across infancy. Finally, our study was not sufficiently powered to examine the role of maternal ART timing on MMc levels in T cell lineage subsets within iHEU. Future studies that address these limitations and build upon our reported findings will be critical to understanding the influence of MMc CD8+ T cells in shaping cellular immunity and susceptibility to infection in both iHEU and iHU.

---

**Supplementary Table 1. Dataset.**

[*Available as download*]

## Footnotes

The authors declare no conflicts of interest.

### Data available in supplementary material

